# GRP78 binds SARS-CoV-2 Spike protein and ACE2 and GRP78 depleting antibody blocks viral entry and infection in vitro

**DOI:** 10.1101/2021.01.20.427368

**Authors:** Anthony J. Carlos, Dat P. Ha, Da-Wei Yeh, Richard Van Krieken, Parkash Gill, Keigo Machida, Amy S. Lee

## Abstract

The severe acute respiratory syndrome coronavirus 2 (SARS-CoV-2), the causative agent of the current COVID-19 global pandemic, utilizes the host receptor angiotensin-converting enzyme 2 (ACE2) for viral entry. However, other host factors may also play major roles in viral infection. Here we report that the stress-inducible molecular chaperone GRP78 can form a complex with the SARS-CoV-2 Spike protein and ACE2 intracellularly and on the cell surface, and that the substrate binding domain of GRP78 is critical for this function. Knock-down of GRP78 by siRNA dramatically reduced cell surface ACE2 expression. Treatment of lung epithelial cells with a humanized monoclonal antibody (hMAb159), selected for its ability to cause GRP78 endocytosis and its safe clinical profile in preclinical models, reduces cell surface ACE2 expression, SARS-CoV-2 Spike-driven viral entry, and significantly inhibits SARS-CoV-2 infection *in vitro*. Our data suggest that GRP78 is an important host auxiliary factor for SARS-CoV-2 entry and infection and a potential target to combat this novel pathogen and other viruses that utilize GRP78.

## Introduction

The coronavirus pandemic caused by the severe acute respiratory syndrome coronavirus 2 (SARS-CoV-2) is currently the greatest threat to global public health. While SARS-CoV-2 vaccines provide optimism to combat COVID-19, identification of targets that may offer therapy for those ineligible for vaccine or infected by escape mutants bypassing vaccine protection is of great interest. While it has been elucidated that the SARS-CoV-2-Spike protein (SARS-2-S) responsible for viral attachment and fusion to the host cells exploits angiotensin-converting enzyme 2 (ACE2) as the cellular receptor for viral entry, evidence is emerging that other host factors may serve as critical entry co-factors for productive infection (1, 2). Recent molecular docking analyses have identified a putative site of interaction between the 78 kilo-Dalton glucose-regulated protein (GRP78) and the receptor binding domain (RBD) of SARS-2-S, raising the interesting possibility that GRP78 can facilitate or serve as an alternative receptor for SARS-CoV-2 entry (3, 4). GRP78, also known as BiP or HSPA5, is broadly expressed in bronchial epithelial cells and the respiratory mucosa at levels significantly higher than ACE2 (5). In recent case-control studies, serum GRP78 levels were found to be elevated in SARS-CoV-2 cases (6). In its capacity as a stress-inducible molecular chaperone, GRP78 is expressed in many cell types and serves critical protein folding functions in the endoplasmic reticulum (ER), and upon translocation from the ER to the cell surface under pathophysiological conditions, acts as co-receptor for various signaling molecules, as well as for viral entry (7–11). For coronaviruses, GRP78 is known to interact with the bat coronavirus HKU9 and MERS-CoV Spike proteins, facilitating cell surface attachment and viral entry (12). Furthermore, virus infection leads to ER stress and increased total and cell surface GRP78 (csGRP78) expression further enhancing viral infection in a positive feedback cycle (11). Here, utilizing biochemical and imaging approaches, we established GRP78 interactions with SARS-2-S and ACE2. We further demonstrated that a humanized monoclonal antibody (hMAb159) with high specificity against GRP78 and a safe clinical profile in preclinical models (13) depletes csGRP78 and reduces cell surface ACE2 (csACE2), SARS-CoV-2 entry and infection.

## Results and Discussion

To test GRP78 binding to SARS-2-S in cells, we expressed HA-tagged SARS-2-S (HA-Spike) and FLAG-tagged GRP78 (F-GRP78) in African green monkey kidney epithelial VeroE6 cells overexpressing ACE2 (VeroE6-ACE2) as a model system. Co-immunoprecipitation (IP) for the HA-epitope showed that F-GRP78 can be pulled down with HA-Spike suggesting potential interaction between the two proteins (Fig. 1*A*). Co-IP with antibody against the FLAG-epitope further showed that F-GRP78 can bind HA-Spike and ACE2 (Fig. 1*A*). Next, we tested the ability of HA-Spike and ACE2 to bind three dominant negative mutants of GRP78: a substrate binding-defective mutant (T453D), an ATP binding-defective mutant (G227D), and a co-chaperone interaction-defective mutant (R197H) (7). Compared to wild type (WT) GRP78, G227D and R197H mutants bound with HA-Spike, albeit at a lower level, while the T453D mutant did not whereas both G227D and T453D mutants were unable to bind ACE2 (Fig. 1*B*). These results indicate that SBD of GRP78 is most critical for interaction between GRP78 and SARS-2-S providing experimental evidence consistent with a previous *in silico* molecular docking study (3), and that GRP78 binding to ACE2 requires both the SBD and the ATP binding domain.

**Figure 1.**
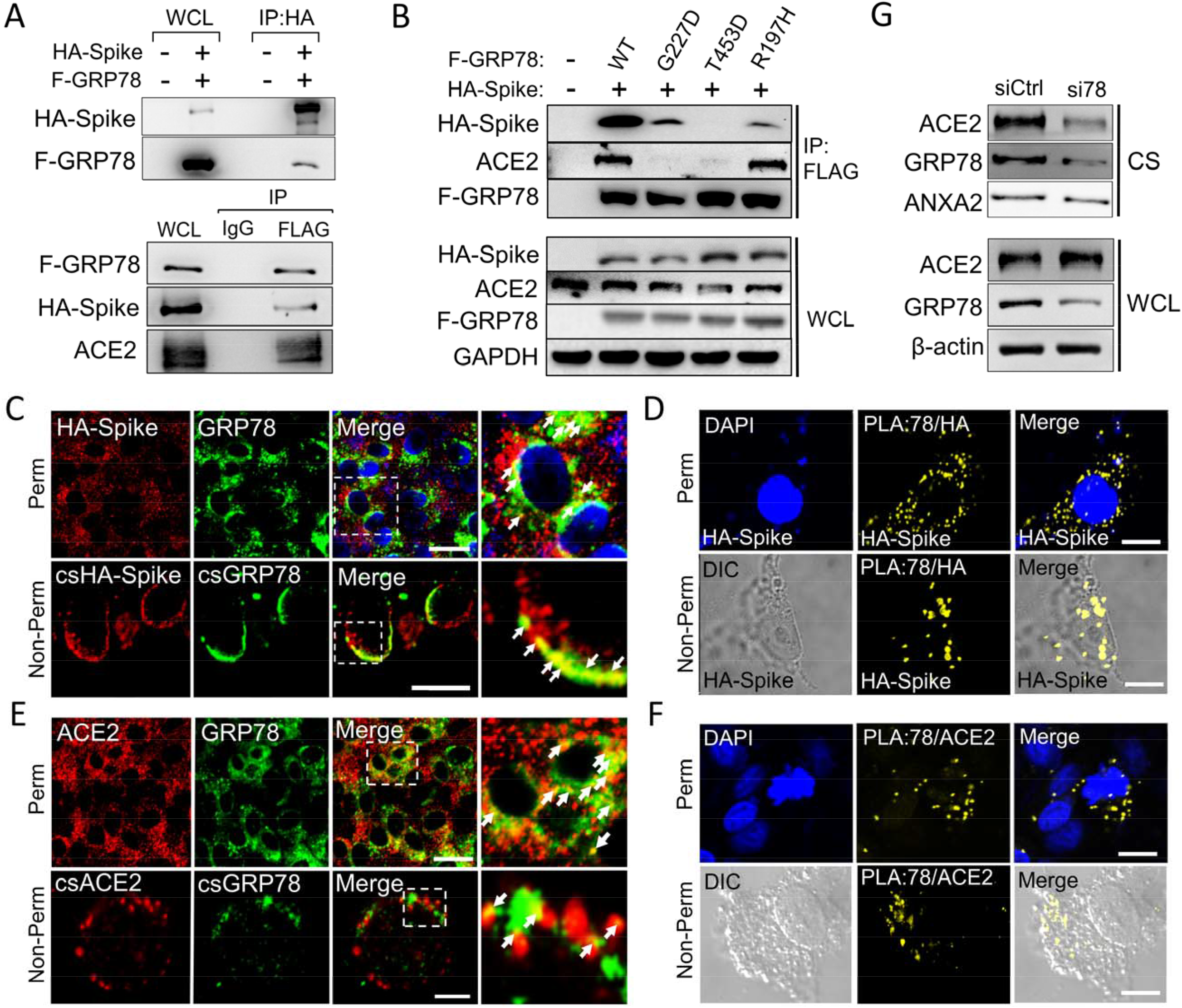
GRP78 interactions with SARS-CoV-2 Spike protein and ACE2. (*A*) Lysates from VeroE6-ACE2 cells expressing FLAG-GRP78 (F-GRP78) and HA-Spike were subjected to immunoprecipitation (IP) using anti-HA, IgG or anti-FLAG antibodies. The whole cell lysate (WCL) and IP fractions were analyzed for the indicated proteins by Western blot. (*B*) Similar to A except lysates from cells expressing wild type (WT) or the indicated mutant forms of GRP78 were subjected to IP with anti-FLAG antibody, with GADPH serving as loading control for the WCL. (*C*) Confocal immunofluorescence (IF) images of VeroE6-ACE2 cells expressing HA-Spike probed with anti-HA (red) and anti-GRP78 (green) antibodies. The top and bottom row represent permeabilized and non-permeabilized cells respectively. The boxed areas are enlarged on the right. Arrows indicate co-staining. (Scale bars, 20 µm) (*D*) VeroE6-ACE2 cells transfected with vector expressing HA-Spike were subjected to proximity ligation assay (PLA) using antibodies against HA and GRP78. DAPI (blue) represents nuclei staining. Yellow indicates colocalization. (Scale bars, 10 µm) (*E*) Similar to C except for IF staining for ACE2 (red) and GRP78 (green). (Scale bars, 20 µm top row, 5 µm bottom row) (*F*) Similar to D except VeroE6-ACE2 cells were subjected to PLA using anti-ACE2 and anti-GRP78 antibodies. (Scale bars, 10 µm) (*G*) WCL or purified cell surface (CS) proteins from VeroE6-ACE2 cells treated with control siRNA (siCtrl) or siRNA against GRP78 (si78) were probed for GRP78 and ACE2 levels by Western blots. β-actin served as loading control for WCL and Annexin 2 (ANXA2) for CS proteins.

Viruses, including SARS-CoV-2, usurp the host ER translational machinery to synthesize the viral proteins in massive quantities. Thus, as a major ER chaperone, GRP78 plays an essential role in viral protein synthesis and maturation (11). Confocal immunofluorescence (IF) microscopy of permeabilized cells expressing HA-Spike showed that it co-localized with endogenous GRP78 in the perinuclear region typical of the ER, and in non-permeabilized cells at the cell surface (Fig. 1*C*). The IF results were confirmed using the Proximity Ligation Assay (PLA) which reveals protein-protein interactions at distances <40 nm (Fig. 1*D*). Likewise, by both confocal IF microscopy and PLA, co-localization between endogenous GRP78 and ACE2 was detected in the perinuclear region typical of the ER, and their co-localization was also observed on the cell surface (Fig.1*E* and *F*). Together, these studies suggest that GRP78 could serve as a foldase for SARS-2-S and ACE2 in the ER, and act as a scaffold for SARS-2-S and ACE2 interaction on the cell surface. Recent studies showed that GRP78 deficiency could lead to decrease in cell surface receptors such as CD109 and CD44 (10, 14). Interestingly, while knockdown of GRP78 by siRNA did not affect total ACE2 protein level, the level of csACE2 decreased markedly in parallel with a decrease in csGRP78 (Fig. 1*G*). Collectively, our results reveal that GRP78, in addition to potentially facilitating SARS-2-S binding to ACE2, may also be important for ACE2 trafficking, localization, and stability on the cell surface, and SARS-2-S production in the ER following viral infection, which awaits future investigation.

To test directly whether csGRP78 facilitates SARS-CoV-2 entry, we employed the human lung epithelial cell line H1299 and the vesicular stomatitis virus (VSV) pseudo particles bearing SARS-2-S as viral entry model system. To specifically target csGRP78, we utilized humanized MAb159 (hMAb159), a monoclonal antibody established to have high specificity and affinity against GRP78 with safe clinical profile in preclinical models (13). In H1299 cells, hMAb159 treatment led to reduced csGRP78 staining (Fig. *2A*), consistent with the ability of MAb159 to cause GRP78 endocytosis and degradation established in other cell systems (13). Flow cytometry analysis of the same cells pretreated with either human IgG1 or hMAb159 showed that hMAb159 reduced both the number of cells expressing csACE2 and the level of csACE2 (Fig. *2B*). In Western blot analysis of H1299 cell lysate, hMAb159 only recognizes a single protein GRP78 and has no cross reactivity with its closely related cytosolic homolog HSP70, reaffirming its high specificity for GRP78 (Fig. 2*C*). In viral entry assays, we observed that pretreatment with hMAb59 significantly reduced SARS-2-S-driven pseudovirus entry at concentration of 0.5 µg/ml but did not affect VSV-G dependent entry into H1299 cells (Fig. *2D* and *E*) or cell viability, which excluded the possibility that the reduced SARS-CoV-2 entry was due to cytotoxicity caused by hMAb159 (Fig. *2F*). Furthermore, VeroE6-ACE2 cells preincubated with hMAb159 prior to infection with live SARS-CoV-2 virus exhibited significant decrease in the number of plaques compared with human IgG1 control (Fig. *2G*). Interestingly, a recent study showed GRP78 co-localizing with SARS-2-S following live virus infection and AR12, an inhibitor of chaperones including GRP78 suppressed SARS-CoV-2 infection (15). Collectively, these results suggest that through targeting host auxiliary chaperones such as GRP78 required for viral entry and production could offer a new strategy toward the development of broad spectrum anti-viral therapy. It is also tempting to speculate that csGRP78 expression elevated in stressed organs may contribute to higher viral entry and morbidity in COVID-19 which warrants further investigation.

**Figure 2.**
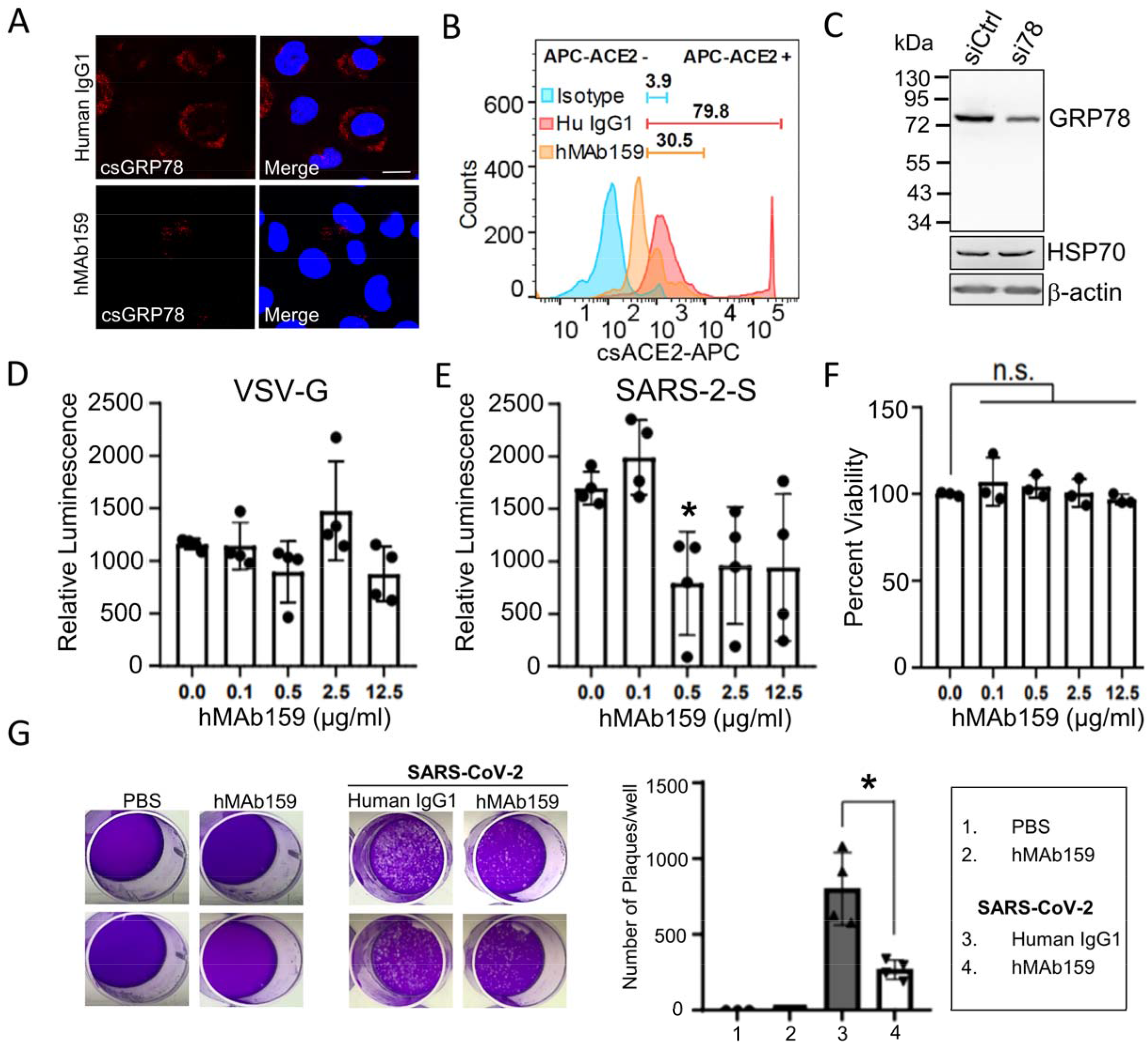
Effect of hMAb159 treatment on cell surface forms of GRP78 and ACE2, viral entry and infection. (*A*) Confocal IF images of non-permeabilized H1299 cells treated with human IgG1 or humanized MAb159 (hMAb159) at 0.5 µg/ml for 2 h at 37°C and probed with anti-GRP78 (red) antibody. DAPI (blue) indicates nuclei staining. (Scale bars, 10 µm). (*B*) Flow cytometry analysis of csACE2 of the same cells treated with human IgG1 or hMAb159. Fluorescence intensity beyond the right border of negative control isotype was set as positive staining. The numbers indicate the percentage of csACE2 positive staining cells under each condition. (*C*) Western blot of H1299 cell lysates treated with control siRNA (siCtrl) or siRNA against GRP78 (si78) and probed with hMAb159 and anti-HSP70 antibody, with β-actin serving as loading control. (*D-E*) H1299 cells pre-incubated with indicated concentration of hMAb159 2 h before transduction were subsequently inoculated with VSV pseudovirus harboring VSV-G (*D*) or SARS-2-S (*E*) surface receptor. At 16 h post-infection, relative luciferase activities were determined. Data are mean ± SD (n = 4). (*F*) In parallel, after 18 h incubation of H1299 with hMAb159 at indicated concentration, cell viabilities were measured via XTT assay. Data are mean ± SD (n = 3). (*G*) Plaque inhibition assay. VeroE6-ACE2 cells pre-incubated with 0.5 µg/ml hMAb159 or human IgG1 for 2 h were subsequently infected with live SARS-CoV-2 at MOI of 0.01. Viral replication was quantified by plaque assay. Data are mean ± SD (n = 4), * denotes p<0.05.

## Methods

Information on the cell lines, biochemical, imaging and viral entry and infection assays are provided in the Supplemental information.

## ACKNOWLEDGEMENTS

This work is supported by NIH grants (R01 CA027607, R01 CA027607-37S1 and R01 CA238029) to A.S.L. and NIH grants (R01 AA025204-01A1, R21 AA025470-01A1 and P50 AA11999) to K. M. We thank Younho Choi for the Vero E6-ACE2 cells, Stefan Pohlmann for the Spike expression plasmids, the Cell and Tissue Imaging Core of the USC Research Center for Liver Diseases for confocal microscopy and the USC Biosafety Lab 3 Core Facility for the plaque assays.

## Notes

### Competing Interest Statement

The authors have declared no competing interest.

