## Extended Materials and Methods for "GRP78 binds SARS-CoV-2 Spike protein and ACE2 and GRP78 depleting antibody blocks viral entry and infection in vitro"

**Supplementary Information**

**Extended Materials and Methods**

**Cell lines and culture conditions**: The VeroE6-ACE2 cell line is a generous gift from Younho Choi at USC. The H1299 cell line was obtained from ATCC. VeroE6-ACE2 cells are cultured in Dulbecco's modified Eagle's medium (DMEM with 4.5 g/l glucose) supplemented with 10% Fetal Bovine Serum (FBS) and 1% Penicillin/Streptomycin and 1μg/ml Puromycin at 37 °C and 5% CO_2_. H1299 cells were cultured in RPMI-1640 supplemented with 10% Fetal Bovine Serum (FBS) and 1% Penicillin/Streptomycin at 37°C and 5% CO_2_.

**Expression Vector Construction and Transfection.** Expression vector for HA-Spike is a generous gift from Stefan Pohlmann, Leibniz Institute for Primate Research, Gottinggen, Germany (1). Wild-type (WT) FLAG GRP78 and GRP78 constructs G227D, T453D, and R197H have been previously described (2). Transfections of plasmids were performed with BioT Transfection Reagent (Bioland Scientific, Paramount, CA) following the manufacturer’s instructions. Transfections of siRNAs were performed using Lipofectamine RNAiMAX Transfection Reagent (Thermo Fisher Scientific, Waltham, MA) according to manufacturer’s instructions. The sequences for the siRNAs are as followed: siCtrl: 5’-GAGAUCGUAUAGCAACGGUdTdT-3’; si78: 5’-GGAGCGCAUUGAUACUAGAdTdT-3’.

**Immunoprecipitation.** Immunoprecipitation was conducted as previously described (3). Briefly, cells were lysed in cell lysis buffer [20 mM Tris·HCl (pH 7.5), 150 mM NaCl, 1 mM EDTA, 1 mM EGTA, 1% Triton X-100] supplemented with Halt protease and phosphatase inhibitor (Life Technologies). The lysate was centrifuged at 14,000 × g at 4 °C for 10 min, and the supernatant was subjected to incubation with Anti-FLAG M2 Affinity Gel (A2220; MilliporeSigma) resin or anti-HA Dynabeads (#88836, ThermoPierce) overnight at 4°C. Samples were eluted at 95°C in Western Blot Sample buffer.

**Cell Surface Biotinylation and Avidin Pull Down.** Biotinylation and pull down of cell-surface proteins were performed as previously described (3).

**Immunoblot Analysis**. Protein samples were subjected to 10% or 12.5% SDS/PAGE and Western blot analysis as previously described (3). The primary antibodies used in this study are as follows: rabbit anti-ACE2 (1:1000; 21115-1-P; Proteintech), mouse anti-HA (1:1000; sc-7392; Santa Cruz Biotechnology, Inc.), mouse anti-FLAG M2 (1:1000; F3165; MilliporeSigma), mouse-anti-GAPDH (1:1000; 365062; Santa Cruz Biotechnology, Inc), mouse anti-HSP70 (1:1000; sc-66048; Santa Cruz Biotechnology, Inc.), mouse anti-β-actin (1:1000; 66009-1-Ig, Proteintech). The secondary antibodies used in this study were HRP-conjugated mouse IgG kappa binding protein (1:1000; sc-516102; Santa Cruz Biotechnology, Inc.), HRP-conjugated mouse anti-rabbit secondary antibody (1:1000; sc-2357; Santa Cruz Biotechnology, Inc.). Protein levels were visualized and quantified by a ChemiDoc XRS+ imager (Bio-Rad Laboratories) and BioRad ImageLab software.

**Flow cytometric analysis**. H1299 cells pre-incubated with either human IgG1 or humanized MAb159 (hMAb159) (4) for 2 h were harvested and resuspended in PBS containing 2% BSA and incubated with rabbit anti-ACE2 antibody (Proteintech Cat# 21115-1-AP) at room temperature for 1 h. Cells were washed and incubated with AlexaFluor 647 conjugated anti-rabbit secondary antibody for 30 min. After washing, cells were analyzed on a FACSVerse flow cytometer (Becton Dickinson, San Jose, CA, USA).

**Immunofluorescent Staining.** Immunofluorescent staining for permeabilized cells was conducted as previously described (3). To visualize cell surface proteins, cells were immediately cooled to 4°C, followed by fixing in cold paraformaldehyde (4^o^C, 4%, 15 min), then blocking buffer using BSA (4^o^C, 5% m/v, 1 h). Primary antibodies diluted in 5% m/v BSA were incubated overnight at 4°C. The primary antibodies used were as follows: rabbit anti–HA-probe (1:100; sc-805; Santa Cruz Biotechnology, Inc.), mouse anti-GRP78 (MAb159) (1:100) (4), rabbit anti-ACE2 (1:100, 21115-1-AP, Proteintech). The secondary antibodies used in this study were Alexa Fluor 488 goat anti-mouse antibody (1:100; A32723; Invitrogen) and Alexa Fluor 647 goat anti-rabbit antibody (1:100; A32733; Invitrogen). Imaging was conducted using a Leica SP8 Confocal system with 63x oil objective and the Leica Application Suite 10 Software package.

**Proximity Ligation Assay.** Duolink Proximity Ligation Assay (PLA) was purchased from Millipore-Sigma (DUO92102, MilliporeSigma). VeroE6-ACE2 cells were plated in a sterile Millicell EZ SLIDE 8-well glass (PEZGS0816, Millipore-Sigma) coated with 5 μg/ml Collagen at a density of 30,000 cells per well. To visualize intracellular colocalization, cells were fixed in paraformaldehyde (4%, 15 min) and subsequently permeabilized in Triton-X (0.1% v/v, 15 min) before starting PLA. Thereafter, the PLA protocol was followed to manufacturer's specifications. To visualize cell surface colocalization, non-permeabilized cells were fixed in paraformaldehyde (4%, 15 min). Thereafter, the PLA protocol was followed to manufacturer's specifications. Imaging was conducted using a Leica SP8 Confocal system with 63x oil objective and the Leica Application Suite 10 Software package. The primary antibodies used are as follows: rabbit anti–HA-probe (1:100; sc-805; Santa Cruz Biotechnology, Inc.), mouse anti-GRP78 (MAb159) (1:100) (4), rabbit anti-ACE2 (1:100, 21115-1-AP, Proteintech). For negative controls, mouse and rabbit IgG isotype-matched antibodies were used.

**Generation of VSV pseudotyped and transduction experiments.** The procedure for VSV pseudoparticles production was following a published protocol (5). Briefly, 293T cells ectopically expressing the viral surface glycoprotein, either VSV-G or SARS-CoV-2-Spike (kindly provided by Stefan Pohlmann, Leibniz Institute for Primate Research, Gottinggen, Germany) were inoculated with a replication-deficient VSV vector, VSVΔG-fLuc (kindly provided by Jae Jung, Department of Molecular Microbiology and Immunology, USC, CA, USA) that expresses firefly luciferase instead of VSV-G. After 1 h incubation at 37 °C, media was removed, and cells were washed twice with PBS before fresh complete media was added. 24 h post-inoculation, media containing pseudotyped particles was harvested and centrifugated. Supernatant clarified from cellular debris was used in transduction experiments. For transduction, H1299 grown to 70% confluency in 96-well plates were inoculated with respective VSV pseudo particles. To block GRP78 on the cell surface, cells were pre-incubated 2 h before transduction with 0.1 to 12.5 μg/ml hMAb159 (4). An unrelated human IgG1 (Bio-Rad, Hercules, CA, USA) was used as control. 16 h post-transduction, the firefly luciferase activity was determined by using ONE-Glo™ Luciferase Assay System (Promega, Medison, WI, USA) and FLUOstar Omega microplate reader (BMG Labtech, Durham, NC, USA).

**Quantification of cell viability.** Cytotoxicity for incubation of H1299 with hMAb159 (4) was measured with the XTT Cell Proliferation Assay Kit (ATCC, Manassas, VA, USA) according to the manufacturer’s instructions. Briefly, H1299 cells grown to 70% confluency in 96-well plate were incubated for 18 h with various concentrations (0.1-12.5 μg/ml) of hMAb159 in 100 μl culture media. Next, 50 μl of activated-XTT solution was added into each well. Wells containing only culture media served as a background control to determine blank absorbance readings. After 4 h, the absorbances at 475 nm and 660 nm were measured using FLUOstar Omega microplate reader (BMG Labtech, Durham, NC, USA). The specific absorbance = A475 nm (Test) – A475 nm (Blank) – A660 nm (Test).

**Plaque Inhibition** **Assay**

VeroE6-ACE2 cells grown to confluence in 24-well plates were pre-incubated with either PBS, isotype matched human IgG1 or hMAb159 (4) for 2 h then incubated with 10-fold serial dilution, 10^-2^ or 10^-3^ of virus at 37°C, 5% CO_2_ shaker for 1 h. After the virus was removed, the cells were washed and added to 1.5 ml agarose/media overlay. After 3 days of incubation at 37°C, VeroE6-ACE2 cells were fixed with 4% formaldehyde for 3 days. After removing formaldehyde and agarose/media plug, the cells were stained with crystal violet and wash with water. The number of plaques was quantified by ImageJ.
